# Two types of motor inhibition after action errors in humans

**DOI:** 10.1101/2022.06.17.496641

**Authors:** Yao Guan, Jan R. Wessel

**Author notes:** ***Corresponding author address:*** Jan R. Wessel, Ph.D., Department of Psychological and Brain Sciences, Department of Neurology, University of Iowa Hospitals and Clinics, 444 MRC, Iowa City, IA 52242, www.wessellab.org.

## Abstract

Adaptive behavior requires the ability to appropriately react to action errors. Post-error slowing of response times (PES) is one of the most reliable phenomena in cognitive neuroscience. It has been proposed that PES is partially achieved through inhibition of the motor system. However, there is no direct evidence for this link – or indeed, that the motor system is physiologically inhibited after errors altogether. Here, we used transcranial magnetic stimulation and electromyography to measure cortico-spinal excitability (CSE) across four experiments using a Simon task, in which human participants sometimes committed errors. Errors were followed by reduced CSE at two different time points, and in two different modes. Shortly after error commission (250ms) CSE was broadly suppressed – i.e., even task-unrelated motor effectors were inhibited. During the preparation of the subsequent response, CSE was specifically reduced at task-related effectors only. This latter effect was directly related to PES, with stronger CSE suppression accompanying greater PES. This suggests that PES is achieved through increased inhibitory control during post-error responses. To provide converging evidence, we then re-analyzed an openly-available EEG dataset that contained both Simon- and Stop-signal tasks using independent component analysis. We found that the same neural source component that indexed action-cancellation in the stop-signal task also showed clear PES-related activity during post-error responses in the Simon task. Together, these findings provide clear evidence that post-error adaptation is partially achieved through motor inhibition. Moreover, inhibition is engaged in two modes (first non-selective, then selective), aligning with recent multi-stage theories of error processing.

## INTRODUCTION

Adjusting performance after erroneous actions is a key component of goal-directed behavior. One long-proposed aspect of error processing is the momentary inhibition of the motor system to increase caution and avoid further errors (Ridderinkhof, 2002; Danielmeier & Ullsperger, 2011). This proposition is supported by some indirect neural evidence. First, BOLD activity in motor cortex decreases after errors (King et al, 2010; Danielmeier et al., 2011). Second, errors evoke neural activity that is also observed during outright action-stopping (Wessel et al., 2012; Marco-Pallarés et al., 2008). Third, errors activate the subthalamic nucleus (Cavanagh et al., 2014; Siegert et al., 2014), which is purportedly key to inhibitory motor control (Aron, 2011; Jahanshahi et al., 2015; Wessel & Aron, 2017). Hitherto, however, there is no direct evidence for the physiological inhibition of the motor system after errors.

According to influential work on action-stopping, the physiological effects of motor inhibition can be either selective (i.e., effector-specific) or non-selective (i.e., cross-effector, Aron 2011). Selective inhibition is limited to the primary responding effector(s) and ostensibly implemented via the indirect pathway of the basal ganglia (Majid et al., 2013). Non-selective inhibition, in contrast, is broad and extends even to task-unrelated effectors (Badry et al., 2009; Wessel & Aron, 2017). It is ostensibly implemented via the hyper-direct pathway (Chen et al., 2020; Wessel et al., 2022a). Empirically, selective and non-selective inhibition can be distinguished through corticospinal excitability (CSE), which can be measured via concurrent transcranial magnetic stimulation (TMS) and electromyography (EMG; Bestmann & Krakauer, 2015; Duque et al., 2017). TMS over the primary motor cortex representation of a muscle produces a motor evoked potential (MEP) in the EMG, whose amplitude reflects the net-CSE of the underlying cortico-spinal tract. In line with the dichotomy between selective and non-selective inhibition, studies using action-stopping paradigms have shown that inhibtion can suppress CSE at task-relevant effectors only (Greenhouse et al., 2012; Majid et al., 2013) or at both task-related and task-unrelated effectors (Badry et al., 2009; Cai et al., 2012), depending on the amount of strategic control. When inhibitory control is rapid and reactive, it is typically non-selectively implemented (Majid et al., 2012; Wessel et al., 2013). However, when it can be deployed strategically and with foreknowledge, it is often selective (Claffey et al, 2010; Cai et al., 2011).

With regards to error processing, the adaptive orienting theory (AOT; Wessel, 2018) hypothesizes that errors trigger both selective and non-selective inhibition at different time periods. Specifically, the AOT suggests that error processing occurs in two sequential steps. The first is a universal, reflexive inhibition-orienting phase, which takes place immediately after errors (and other expectancy violations). During this phase, behavior and cognition are purportedly transiently inhibited, which facilitates an orienting process towards the source of the expectancy violation (Corbetta et al., 2008; Dayan & Yu, 2006; Maier et al., 2011). Once the action error is identified as that source, a strategic, error-specific phase begins (Danielmeier et al., 2011; Ullsperger, & Von Cramon, 2001; Purcell & Kiani, 2016; Cavanagh et al., 2011; Gjorgieva & Egner, 2022). According to the AOT, the motor system will be broadly and non-selective inhibited during the initial inhibition-orienting stage, whereas inhibition in the subsequent strategic phase will be effector-specific.

We tested these hypotheses in 5 experiments using the Simon task (Simon & Rudell, 1967). In Experiments 1-3, we measured CSE at task-unrelated effectors immediately after response commission (i.e., during the post-error inhibition-orienting phase). We predicted that CSE would be non-selectively suppressed after errors compared to correct responses. In Experiment 4, we measured CSE after the presentation of the following stimulus (i.e., in the strategic adjustment phase during which the next response is prepared). We predicted that on post-error trials, CSE would be selectively suppressed at the task-relevant effectors, and that this suppression would relate to post-error behavioral adjustments. In Experiment 5, we then used EEG to buttress these experiments by showing that a functionally-localized neural signature of selective motor inhibition is also involved in adjusting behavior during the strategic post-error period.

## MATERIALS AND METHODS

### Participants

In Experiments 1 and 2, we recruited 57 right-handed college students (*N*_Exp1_ = 42, *M*_age_ = 20.48 years, *SD*_age_ = 2.94, 16 males; *N*_Exp2_ = 15, *M*_age_ = 21.73 years, *SD*_age_ = 6.71, 7 males) in exchange for course credit. In Experiments 3 and 4, we recruited 111 right-handed college students (*N*_Exp3_ = 76, *M*_age_ = 19.07 years, *SD*_age_ = 2.68, 20 males; *N*_Exp4_ = 35, *M*_age_ = 19.83 years, *SD*_age_ = 2.87, 8 males) in exchange for course credit or an hourly payment of $15. For Experiment 5, we re-analyzed an existing EEG dataset (Experiment 1 from Wessel, Waller, & Greenlee, 2019) with 21 healthy adult participants (*M*_age_ = 27.40 years, *SEM*_age_ = 2.24, 10 males). Written informed consent was collected before the experiment. All procedures were approved by the local Institutional Review Board and conducted in accordance with the Declaration of Helsinki.

### Experimental tasks

Stimuli were presented using PsychToolbox (Brainard, 1997) in MATLAB (R2017a) via a Ubuntu Linux desktop computer. We used the same visual Simon task (***Figure 1***) from Wessel et al. (2019) across all experiments. Each trial consisted of a 500 ms fixation, followed by a 200 ms colored square (target), presented either to the left or right of the fixation cross. The color of the square indicated the response. The color-response mappings were displayed on the bottom of the screen throughout the experiment. The response window (up to 1500 ms) was followed by an inter-trial interval of 1000 ms. Participants completed a total of 768 trials across 8 blocks of 96 trials each after a practice session. Half of the trials were congruent (the side of the response matched the side of the stimulus presentation), whereas the other half was incongruent. In between blocks, participants received feedback about their performance (i.e., reaction time, number and percentage of response errors and misses). In Experiments 1-3 (TMS), participants responded with their feet using foot pedals. In Experiment 4 (TMS), participants responded with index and middle finger from the right hand. In Experiment 5 (EEG), participants responded with both hands.

**Figure 1.**
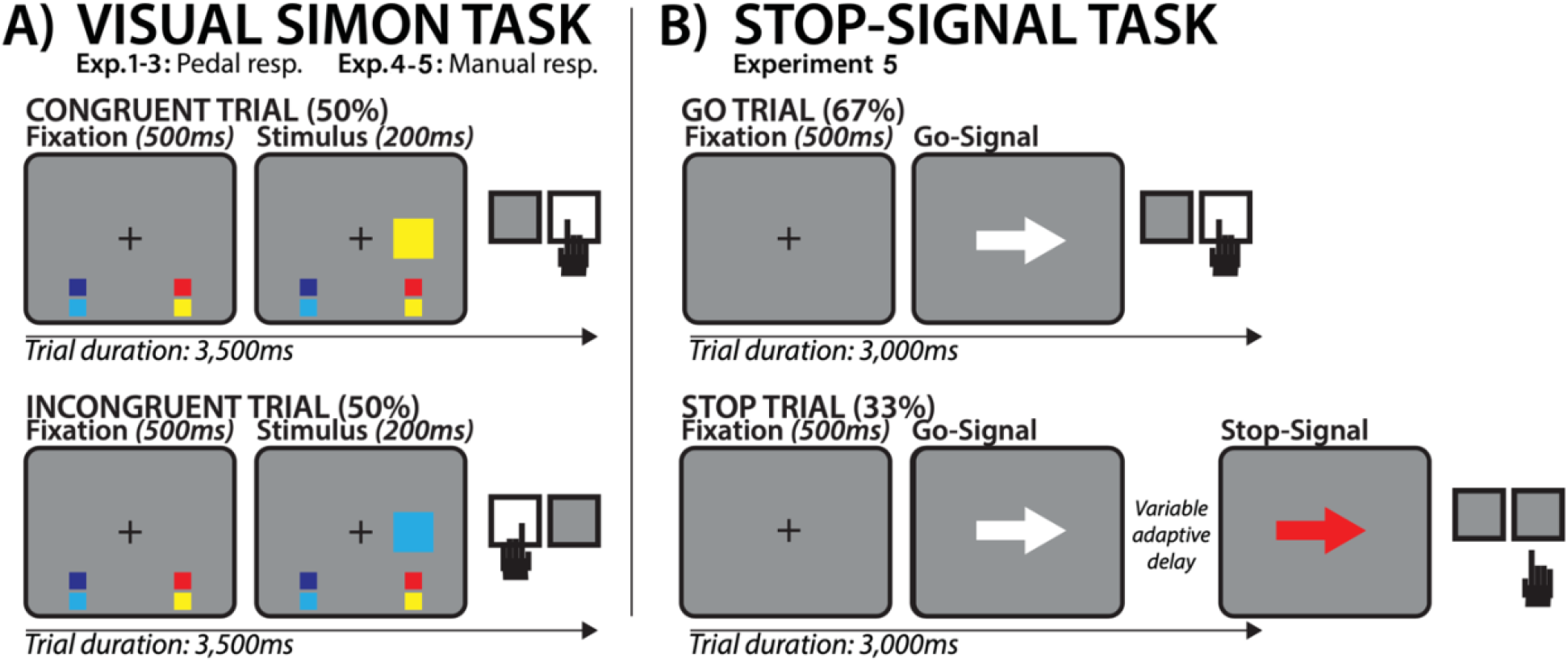
Diagrams of experimental paradigms. Left: Visual Simon task (Experiments 1-5) and visual stop-signal task (Experiment 5). Figure reproduced from Wessel et al., 2019.

Experiment 5 (EEG) also used data from a stop-signal task, collected in the same session after the visual Simon task, described in Wessel et al. (2019). A full description can be found therein. Each trial consisted of a fixation and a go-signal. Participants were instructed to respond to the direction of the go-signal. On 33% of the trials, a stop-signal occurred following the go-signal at variable adaptive delay. Participants needed to withhold their manual response in response to the stop-signal.

### EMG recordings (Experiments 1-4)

In Experiments 1-3, EMG was measured using a bipolar belly-tendon montage from the first dorsal interosseus (FDI) muscle of the right hand using adhesive electrodes and a ground electrode placed over the distal end of ulna. In these three experiments, participants responded using their feet, rendering the FDI muscle task-unrelated. In Experiment 4, EMG was recorded from the right flexor digitorum superficialis (FDS), with a ground electrode placed over the distal end of the shoulder clavicle bone. The FDS controls the flexion of the middle phalanges of the hand, which were used to make responses in Experiment 4, rendering the FDS muscle task-related.

### TMS stimulation (Experiments 1-4)

TMS was performed as described in Dutra, Waller, & Wessel (2018), the description is adapted from there. TMS stimulation was delivered using a MagStim 200-2 system with a 70 mm figure-eight coil. Hotspotting was conducted to identify the FDI (Experiments 1-3) or FDS (Experiment 4) stimulation locus and intensity. The coil was first placed 5 cm lateral and 2 cm anterior to the vertex and repositioned to where the largest MEPs were observed consistently. Resting motor threshold (RMT) was then defined as the minimum intensity required to induce MEPs of amplitudes exceeding 0.1 mV peak to peak in 5 to 10 consecutive probes (Rossini et al., 1994). TMS stimulation intensity was then adjusted to 115% of RMT for stimulation during the experimental task. An EMG sweep was started 100 ms before the TMS stimulation.

In Experiment 1, TMS stimulation was delivered immediately after each response, at a delay of either 150, 200, or 250ms (in a group of *N* = 20) or 300, 375, or 450ms (in a separate group of *N* = 19).

Since we found significant error-related CSE suppression at 250ms after errors in Experiment 1, we replicated this result in Experiment 2 using only that time point (i.e., CSE was collected at 250ms after every response) in a group of *N* = 15. The sample size in Experiment 2 was based on a power calculation from the effect size at the 250ms time point in Experiment 1.

Experiment 3 used the same time points as Experiment 1 (150, 200, 250ms in a group of *N* = 31 and 300, 375, 450ms in a group of *N* = 45, except that these times refer to stimulus onset rather than response onset. The sample size in Experiment 3 was larger than the sample size in Experiment 1, since we expected a null finding and wanted to ensure sufficient power.

In Experiment 4, CSE was collected at 200, 250, or 300ms after stimulus onset in *N* = 35. The sample size in this experiment was determined based on our hypothesis of a cross-subject correlation between selective post-error CSE suppression and post-error slowing.

### TMS data preprocessing and motor evoked potentials analysis (Experiments 1-4)

TMS data were preprocessed to quantify the MEP. Trials were excluded if they met one of the following criteria: 1) Noisy baseline in EMG recording (root mean square of pre-pulse baseline period exceeded .1 mV); 2) EMG signal exceeded +/- 2.99 mV; 3) No MEP (MEP amplitude < .05 mV). The MEP amplitude was quantified with a peak-to-peak rationale, measuring the difference between maximum and minimum amplitude within a time period of 10 – 40 ms after the pulse. MEP amplitudes were then averaged for each condition of interest and normalized by dividing the mean of the congruent correct trials, block by block and stimulation timing by stimulation timing for each participant.

### In- and exclusion (Experiments 1-4)

Participants were excluded due to 1) procedural issues (e.g., premature termination of the experiment, hardware malfunction); 2) insufficient trial count in one of the experimental conditions of interest (typically in one of the time points of the error condition; minimum cutoff for inclusion: 7 trials); 3) outliers in conditional MEP amplitude (Grubbs test at *p* < .05); and 4) noisy EMG / no reliable MEP.

In Experiment 1, we excluded *N* = 3 for procedural issues, *N* = 3 for outliers in the MEP, and *N* = 2 for noisy EMG, leaving *N* = 34 across two groups (150, 200, 250 ms group: *N* = 17, *M*_age_ = 19.18 years, *SD*_age_ = 1.33, 8 males; 300, 375, 450 ms group: *N* = 17, *M*_age_ = 20.88, years, *SD*_age_ = 2.02, 6 males).

In Experiment 2, we excluded *N* = 1 for outliers in the MEP, leaving *N* = 14 (*M*_age_ = 21.86 years, *SD*_age_ = 6.95, 7 males).

In Experiment 3, we excluded *N* = 4 for procedural issues, *N* = 20 for insufficient trials, and *N* = 2 for outliers in the MEP, leaving *N* = 50 across two groups (150, 200, 250 ms group: *N* = 20, *M*_age_ = 19.90, years, *SD*_age_ = 4.99, 7 males; 300, 375, 450 ms group: *N* = 30, *M*_age_ = 18.70 years, *SD*_age_ = 0.75, 6 males).

In Experiment 4, we excluded *N* = 2 for procedural issues, *N* = 1 for outliers in the MEP, and *N* = 1 for noisy EMG, leaving *N* = 31 (*M*_age_ = 20 years, *SD*_age_ = 3.01, 7 males).

### Statistical analyses of behavioral data and MEP amplitudes (Experiments 1-4)

To quantify the response conflict effect in the Simon task, we conducted paired-samples *t*-tests to compare reaction time and error rate in correct incongruent vs. correct congruent trials, respectively. To quantify post-error slowing (PES), we conducted paired *t*-test to compare reaction time in correct trials following incongruent error vs. incongruent correct trials.

We then examined how MEP amplitudes changed as a function of trial accuracy (error or correct in Experiments 1 and 2; pretrial-error or pretrial-correct in Experiments 3 and 4) and TMS stimulation timing (150, 200, 250, 300, 375, or 450 ms in Experiment 1 and 3; 200, 250 or 300 ms in Experiment 4) with respect to the critical event (response or stimulus). To account for the data structure (the six stimulation time points in Experiments 1 and 3 were realized across two separate groups of participants) and individual differences (Yu et al., 2021), we conducted linear mixed-effects modeling (Boisgontier & Cheval, 2016; Volpert-Esmond et al., 2018) with crossed random effect for subjects (Judd et al., 2012) in Experiments 1, 3, and 4. Specifically, we fit mixed-effects models for trial-level TMS data using the R package *lme4* (Bates et al., 2015). The dependent variable was the trial-level normalized MEP amplitude. We applied a natural log transformation to the normalized MEP amplitude because its residual distribution was right-skewed. The mixed variables were trial accuracy and TMS stimulation timing. Subjects were included as a random variable, with intercepts being random effects. We focused on the ANOVA results from the mixed-effects models to test the omnibus effect of the predictors.

Experiment 2 was a replication of Experiment 1 and only included a single time point (250 ms). Here, we conducted a one-sided paired *t*-test because we had had a directional a priori hypothesis from Experiment 1.

Additionally, in Experiment 4 we tested the cross-subject correlation between the amount of MEP suppression on error (compared with correct) trials and the amount of reaction time slowing. To do so, we fit the subject-level reaction time data into a robust linear regression model with MEP amplitude as the predictor. Both the MEP change and the reaction time change between errors and correct trials were quantified in terms of % change.

### EEG data processing (Experiment 5)

Preprocessed EEG data were taken from Wessel et al., (2019), a publicly available dataset (https://osf.io/k3ypt). Data were preprocessed as described therein. In that study, the combined EEG recordings from the Simon- and stop-signal tasks were preprocessed jointly and subjected to a common infomax independent component analysis (ICA). The stop-signal portion of each subject’s data were then used to functionally localized the neural source (i.e., independent component) that contained the stop-signal P3 event-related potential (ERP). In the stop-signal task, the P3 component shows an earlier onset on successful vs. failed stop-trials and correlates with stop-signal reaction time. These features were quantified to validate the selection of the component from the stop-signal portion of the data (both pictured in Figure 4B, which was reproduced from Wessel et al., 2019). Moreover, their time-frequency decomposition shows a dominance of lower frequencies (delta and theta; see Figure 4A), in line with prior work on time-frequency decompositions of stop-signal task data (Huster et al., 2013). In Wessel et al., (2019), the activity of this component was then interrogated in the Simon task portion of the EEG recording, specifically to investigate if there were significant differences in its activity on incongruent correct vs. congruent correct trials. Error trials were not analyzed in that study.

Here, we applied the same analysis logic (using the same functionally localized stop-signal P3 components from the stop-signal task portion of each participant’s data) and interrogated the activity of these components on correct post-error trials. Based on the results from TMS Experiments 3 and 4 of the current study, we predicted that the independent component that explained the stop-signal P3 would show activity during that same time period and relate to PES (as the stop-signal P3 has recently been proposed to reflect a selective, specific inhibitory process that purportedly underlies selective CSE inhibition, c.f., Diesburg & Wessel, 2021).

To this end, we reconstructed the EEG signal from the Simon task portion of each recording using only the independent component that was identified as underlying the stop-signal P3 in the stop-signal portion. We then constructed a time-frequency decomposition using the filter-Hilbert method, as described in Wessel et al., (2019). Power values were extracted by squaring the magnitude of the resulting complex signal at each of the 30 frequencies of interest (linearly spaced from 1-30Hz), which were then epoched from 300 ms prior to 700 ms post-stimulus. Baseline correction was implemented by subtracting the mean amplitude of each frequency in the baseline window (−300 to 0) from each time-frequency point and dividing by the standard deviation of the amplitude of its frequency in the time range of the baseline window. We then applied a single-trial level generalized linear model (GLM) to the stop-signal P3 components in the EEG signal of the Simon task portion. Specifically, we interrogated activity on correct trials that followed errors. We regressed the *z*-score of the reaction time on those trials onto the full-spectrum time-frequency power from the EEG. The *t* statistics was subsequently used to test for significant differences from 0 on the group-level. The false-discovery rate procedure (Benjamini & Hochberg, 1995) was used for multiple comparison correction with the desired false discovery rate being .05.

## RESULTS

### Behavior

Behavioral results from the Simon task across the 5 experiments are shown in ***Table 1*** (Experiment 5 results are reproduced from Wessel et al., 2019). Correct incongruent trial reaction times were slower than correct congruent trial reaction times, and percent accuracy on correct incongruent trials was lower than correct congruent trial, indicating response conflict. Post-error trial reaction times were slower than post-correct trial reaction times, indicating significant PES.

**Table 1.** Behavioral results in Experiments 1-5. We compared correct incongruent vs. correct congruent trials in RT and error rate, respectively, in the Simon task across 5 experiments. We also compared reaction time in correct trials following incongruent error vs. incongruent correct trials in the Simon Task in Experiments 1-4 (PES). In Experiment 5, we compared correct-trial reaction time with failed-stop trial reaction time in the Stop-signal Task (SST). Note: Experiment 5 results are reproduced from Wessel et al., 2019.

|  | Experiment 1 | Experiment 2 | Experiment 3 | Experiment 4 | Experiment 5 |
| --- | --- | --- | --- | --- | --- |
| **RT** | 469ms [SEM: 7] | 485ms [SEM: 15] | 512ms [SEM: 5] | 422ms [SEM: 6] | 361ms [SEM: 8] |
|  | vs. | vs. | vs. | vs. | vs. |
|  | 431ms [SEM: 7] | 451ms [SEM: 12] | 467ms [SEM: 5] | 384ms [SEM: 6] | 326ms [SEM: 7] |
| | $t(33) = 12.63,$ | $t(13) = 5.61,$ | $t(49) = 16.52,$ | $t(30) = 16.86,$ | $t(20) = 9.63,$ |
| | $p < .001,$ | $p < .001,$ | $p < .001,$ | $p < .001,$ | $p < .001,$ |
| | $d = 2.17.$ | $d = 1.50.$ | $d = 2.34.$ | $d = 3.03.$ | $d = 1.06.$ |
| **Error** | 13.75% vs. 6.4% | 11.80% vs. 5.87% | 15.93% vs. 7.61% | 18.46% vs. 8.07% | 22.67% vs. 10.38% |
| **rate** | $t(33) = 6.29,$ | $t(12) = 3.58,$ | $t(48) = 10.72,$ | $t(30) = 9.44,$ | $t(20) = 4.6,$ |
| | $p < .001,$ | $p = .004,$ | $p < .001,$ | $p < .001,$ | $p < .001,$ |
| | $d = 1.08.$ | $d = .99.$ | $d = 1.53.$ | $d = 1.70.$ | $d = 1.4.$ |
| **PES /** | 472ms [SEM: 9] | 483ms [SEM: 15] | 516ms [SEM: 7] | 430ms [SEM: 8] | 551ms |
| **SST** | vs. | vs. | vs. | vs. | vs. |
|  | 449ms [SEM: 7] | 469ms [SEM: 14] | 486ms [SEM: 5] | 400ms [SEM: 6] | 469ms |
| | $t(33) = 4.23,$ | $t(13) = 2.01,$ | $t(49) = 6.24,$ | $t(30) = 7.57,$ | $t(20) = 14.45,$ |
| | $p < .001,$ | $p = .065,$ | $p < .001,$ | $p < .001,$ | $p < .001,$ |
| | $d = .73.$ | $d = .54.$ | $d = .88.$ | $d = 1.36.$ | $d = 1.44.$ |

### Cortico-spinal excitability

#### Experiment 1

Experiment 1 tested whether errors were followed by a non-selective suppression of CSE at task-unrelated effectors. CSE was measured at six time points (split across two groups of participants) following the response. Mixed-effects ANOVA (***Table 2***) showed an interaction between trial accuracy and TMS stimulation timing, *F*(5, 12013.1) = 2.45, *p* = .031 (***Figure 2A***). Post-hoc paired *t*-test showed that this was explained by the presence of non-selective CSE suppression at 250ms after action errors versus correct responses, *t*(16) = 2.40, *p* = .029, *d* = .58.

**Table 2.**
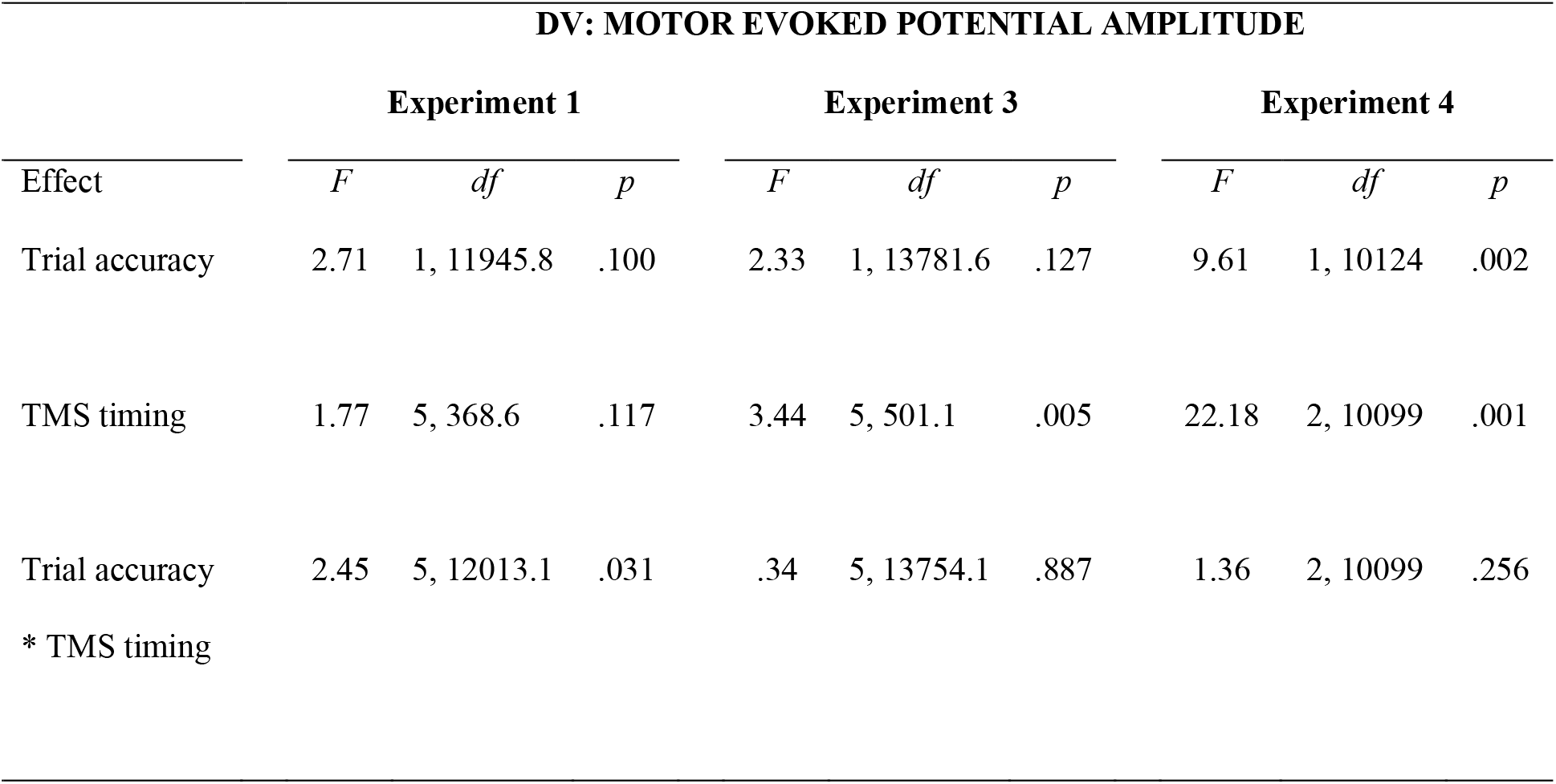
Mixed-effects AVOVA results for MEP in Experiments 1, 3, and 4. MEP amplitudes as a function of trial accuracy (error vs. correct in Experiment 1; pretrial-error vs. pretrial-correct in Experiments 3 and 4) and TMS stimulation timing with respect to the critical event (response or stimulus; 150, 200, 250, 300, 375, or 450 ms in Experiments 1 and 3; 200, 250 or 300 ms in Experiment 4) in the mixed-effects ANOVA.

**Figure 2.**
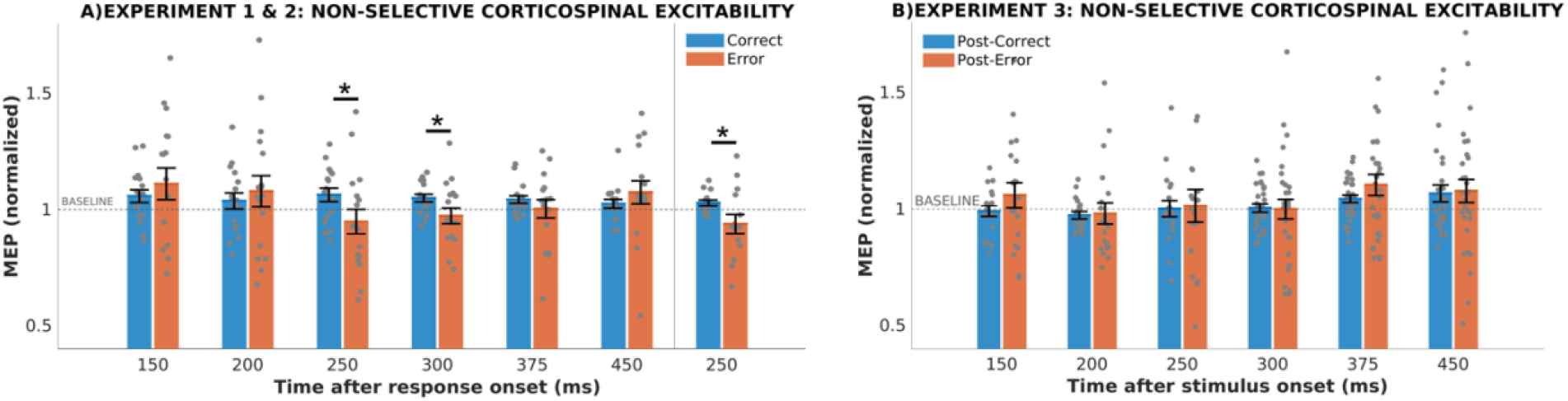
Motor evoked potentials results from the Simon task in Experiment 1-3. Experiment 1 showed significant non-selective motor inhibition at 250 ms after error responses compared to correct responses. Experiment 2 replicated this finding at 250 ms (i.e., during the inhibition-orienting phase after errors). Experiment 3 showed that non-selective motor inhibition did not occur when TMS stimulation timings were stimulus-locked (i.e., during the strategic adjustment phase after errors). Note. *** p < .001, ** p < .01, * p < .05.

#### Experiment 2

Experiment 2 aimed to replicate the finding from Experiment 1 using only the 250ms time point, at which the CSE suppression was significant in Experiment 1. Indeed, a significant suppression on errors was found again, *t*(13) = 1.904, *p* = .04, *d* = .51. Together, Experiments 1 and 2 suggest that CSE is non-selectively suppressed at 250ms after errors compared to correct responses, supporting our hypothesis of non-selective motor inhibition during the inhibition-orienting phase after errors.

#### Experiment 3

Experiment 3 tested whether there was any non-selective inhibition of CSE in the purported strategic period after errors, i.e., after stimulus presentation on correct post-error trials. Mixed-effects ANOVA showed a main effect of TMS timing (***Figure 2B***), *F*(5, 501.1) = 3.44, *p* = .005 (***Table 2***), but no main effect of trial accuracy (*p* = .127) and no interaction (*p* = .887). None of the six time points showed any significant differences between errors and correct responses, with all *p*s > .195 (uncorrected). The time point with the lowest *p* value (375ms) still showed Bayes Factor evidence towards the null (BF_01_ = 2.33). These findings indicate that there was no non-selective CSE suppression during the strategic adjustment phase after errors.

#### Experiment 4

Experiment 4 was designed to test whether the same strategic post-error period that was investigated in Experiment 3 would show selective inhibition at the task-relevant muscle, in line with a purportedly strategically deployed inhibition (***Figure 3***). It was identical to Experiment 3, except that CSE was collected from the responding muscle, not a task-irrelevant one. Indeed, mixed-effects ANOVA showed a main effect of trial accuracy, *F*(1, 10124) = 9.61, *p* = .002, and a main effect of TMS stimulation timing, *F*(2, 10099) = 22.18, *p* = .001 (***Table 2***), with no interaction. Post-hoc paired *t*-test showed MEP suppression in post-error vs. post-correct trials at 200 ms, *t*(30) = 2.98, *p* = .006, *d* = .54, and 250 ms, *t*(30) = 2.21, *p* = .035, *d* = .40.

**Figure 3.**
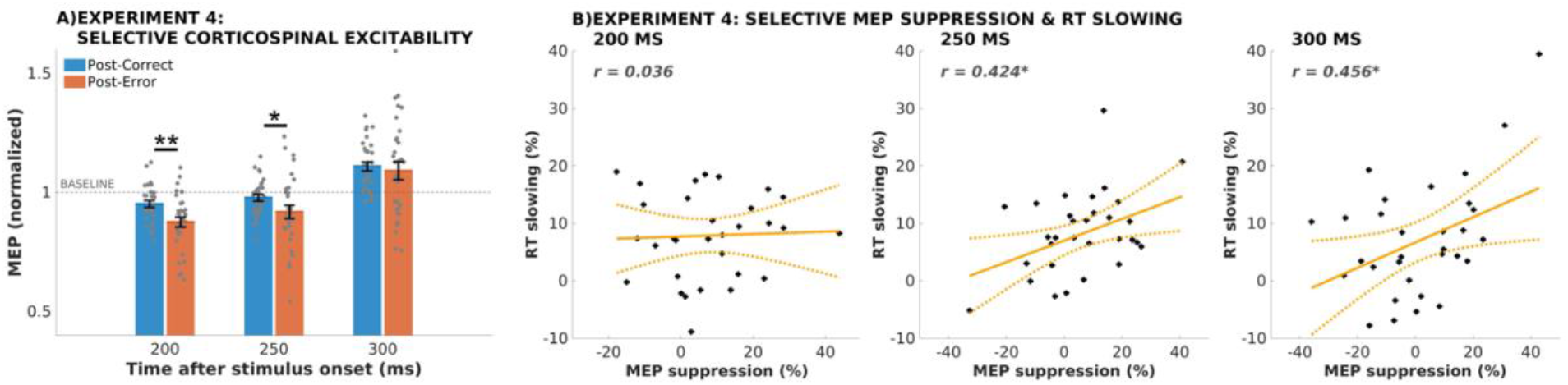
Motor evoked potentials results from Experiment 4. A) Stimulus-locked MEP results showed that CSE was selectively suppressed at 200 and 250 ms in post-error vs. post-correct trials (i.e., during the strategic adjustment phase after errors). B) Correlation results showed at 250 ms & 300 ms after stimulus presentation, the amount of MEP suppression on post-error (vs. post-correct) trials and the amount of RT slowing on post-error trials were positively correlated.

To ensure that these differences in mean CSE could not be explained by differences in raw reaction time between post-error and post-correct trials, we conducted a control analysis that only included a subset of post-correct trials that matched the reaction time from a post-error trial at each TMS stimulation timing. The mixed-effects ANOVA still showed a main effect of trial accuracy, *F*(1, 3298.7) = 4.04, *p* = .045, and a main effect of TMS stimulation timing, *F*(2, 3309.7) = 14.85, *p* < .001, with no interaction, *F*(2, 3298.6) = .43, *p* = .653.These results suggest that errors lead to a selective suppression of CSE at the task-relevant muscle during the preparation of the next response. These findings support our hypothesis of selective motor inhibition during the strategic adjustment phase after errors.

Finally, a cross-subject correlation showed a positive correlation between the relative amount of MEP suppression on error (compared with correct) trials and the amount of relative RT slowing on errors (compared to correct trials, i.e., PES) at 250 ms, *r* = .437, *p* = .012, and 300 ms, *r* = .438 *p* = .012 (MEP and reaction time deltas between errors and correct trials were quantified in terms of % change). This indicates that subjects that show stronger deployment of strategic, selective motor inhibition during the preparation of the next response after an error also show greater post-error slowing. To our knowledge, this is the first demonstration of the fact that post-error slowing is directly related to the physiological suppression of the motor system after errors.

### Source-level EEG (Experiment 5)

The selected independent components reflected the known properties of the P3 in the stop-signal task, as reported in Wessel et al., (2019). In short, they showed an earlier onset on successful compared to failed stop-trials and their onset across subjects was positively correlated with stop-signal reaction time (SSRT) – i.e., subjects with slower SSRT also showed slower P3 onset latencies. Furthermore, the time-frequency decomposition of the components showed that it was dominated by activity in the theta and delta frequencies (***Figures 4A***,***B***).

**Figure 4.**
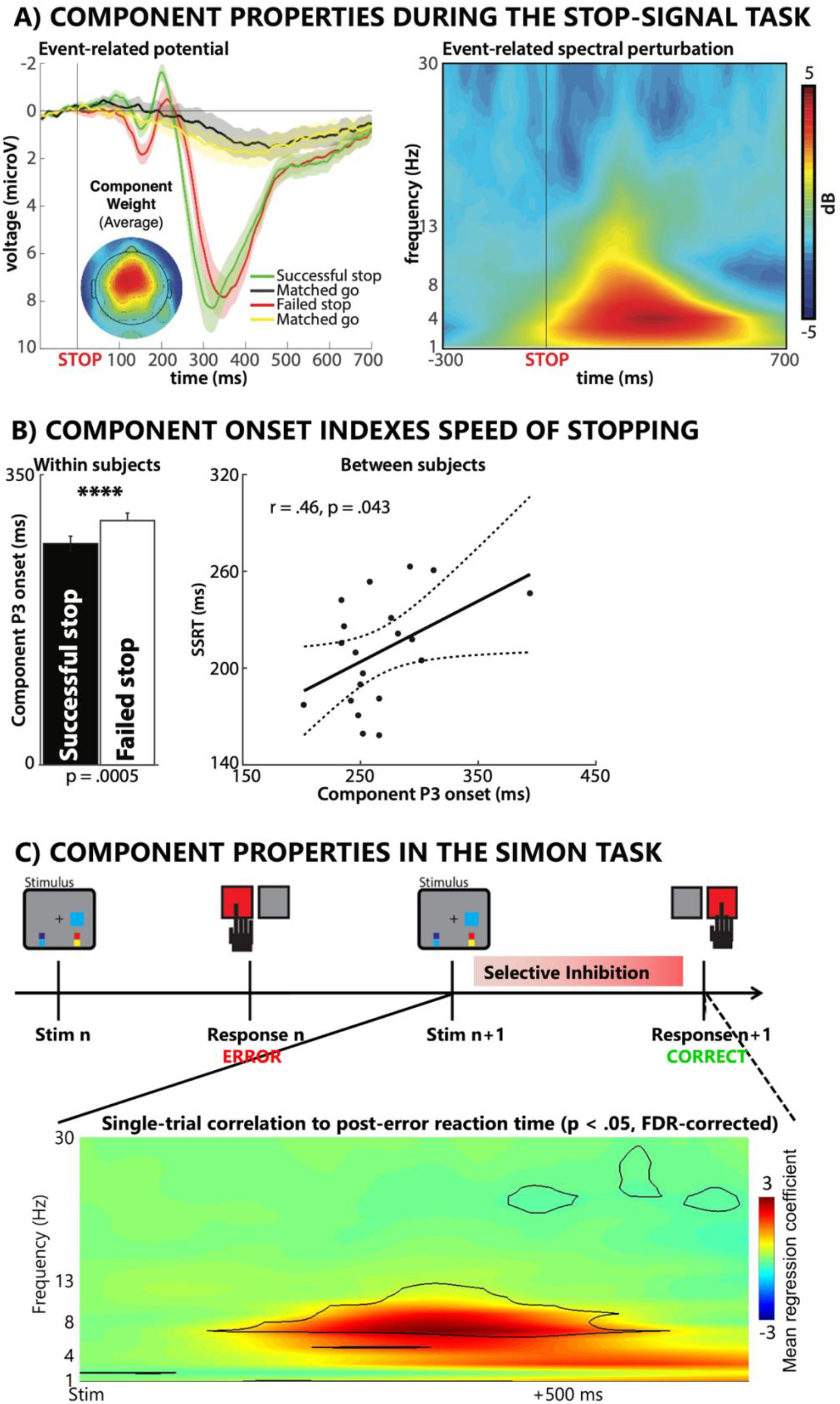
EEG results from Experiment 5. A) Group averaged stop-signal P3 event-related potential and event-related spectral perturbation of the selected independent EEG source-signal components during the stop-signal task. Reproduced from Wessel et al. (2019). B) Onset of the stop-signal P3 shown in panel A positively correlated with the speed of stopping (SSRT) across subjects. Reproduced from Wessel et al. (2019). C) t statistics for regression coefficients from a single-trial level GLM for each time-frequency point after post-error stimuli. The black contour delineates significant group-level deviations of the regression coefficients from 0 (FDR-corrected). The results showed stop-signal P3 components positively correlated with PES in the low frequency during the strategic adjustment phase after errors in the Simon task.

Investigating the activity of the same component in the Simon task, we found that as predicted, its stimulus-locked component activity on post-error trials was positively correlated with reaction time, specifically in the lower frequencies (p < .05, FDR-corrected; ***Figure 4C***). This suggests that in the same post-error time period during which CSE was selectively suppressed in a manner proportional to PES (cf., Experiment 4), the neural source component that indexes inhibitory control in the stop-signal task also showed a direct proportional relationship to PES.

## DISCUSSION

In the current study, we provide first evidence that action errors are followed by a physiological inhibition of the motor system. Moreover, we found that there are two types of motor inhibition that accompany error processing. First, a broad, non-selective suppression that affects even task-unrelated muscles and takes places shortly (around 250ms) after error commission (Experiments 1 and 2), followed by a specific, selective suppression that affects only task-related muscles and takes places during the preparation of the next response (Experiments 3 and 4). These findings provide a key physiological observation that supports the purported role of inhibitory control in error processing – a hypothesis that goes back to at least Burns (1971), but also is key to many subsequent theories of error processing (Botvinick et al., 2001; Ridderinkhof, 2002; Danielmeier & Ullsperger, 2011; Wessel, 2018).

Indeed, it has long been proposed that inhibition is one of the key control processes involved in adapting behavior after action errors. Post-error slowing, the increase of correct-trial reaction time that is observed after error commission, is one of the most reliable findings in the domain of cognitive control (Rabbitt & Rodgers, 1977; Laming, 1979; Notebaert et al., 2009; Dutilh et al., 2012). One of the earliest scholars to propose a role for inhibitory motor processes in the generation of post-error slowing was Burns (1971). Ridderinkhof (2002) later expanded on this possibility using sophisticated behavioral analyses. However, ultimately, behavior alone cannot provide direct evidence for the presence of inhibition, as the slowing of reaction time can – in principle – be caused by a myriad of possible processes that do not necessarily involve inhibitory control (e.g., attentional orienting). As mentioned in the introduction, some preliminary correlational evidence for the presence of motor inhibition after errors comes from reduced BOLD activity in motor cortex after errors (driven by the purported performance-monitoring areas in the medial wall, King et al., 2010). Moreover, the increased activity of the subthalamic nucleus after errors in both time periods of interest in the current study – i.e., both immediately after error commission (Siegert et al., 2014) and following post-error stimuli (Cavanagh et al., 2014) – strongly implies the presence of inhibition. The subthalamic nucleus is a phylogenetically old structure, which is at the core of most neuroscientific models of motor control (Jahanshahi et al., 2015; Frank 2007). It is part of both the indirect and hyperdirect basal ganglia pathways that ostensibly implement both selective and non-selective inhibition of the motor system via excitatory projections to the output nuclei of the basal ganglia (Parent & Hazrati, 1995; Mink, 1996; Wessel & Aron, 2017). Indeed, in two previous studies using the stop-signal task, we have found that the activity of the subthalamic nucleus after stop-signals is directly related to the observed reductions in CSE (Wessel et al., 2016a; Wessel et al., 2022a). However, since the subthalamic is purported to be involved in other, potentially non-motor functions, especially during cognitive control (e.g., Cavanagh et al., 2011; Aron et al., 2016; Zénon et al. 2016; Weintraub & Zaghloul 2013; Temel et al., 2005), its activity alone does not definitively prove the presence of motor inhibition during error processing either. Therefore, we have here used a direct, physiological index of motor system excitability, and have found that errors indeed trigger a physiological inhibition of the motor system, as measured by CSE.

Moreover, the current study goes beyond a mere demonstration of the physiological inhibition of the motor system after errors. Specifically, we show that, in line with the adaptive orienting theory of error processing (Wessel, 2018), errors trigger both selective and non-selective inhibition. The AOT predicts two consecutive steps for error processing, the first of which (inhibition-orienting) purportedly includes non-selective inhibition (see Box 1 in Wessel, 2018). That prediction is based on the assumption that the inhibition-orienting phase is not specific to action errors, but occurs after any type of salient event. Indeed, past work has shown that errors and error-unrelated unexpected events activate an overlapping network of brain regions (Wessel et al., 2012; 2014; Alexander & Brown, 2011) and that expectancy modulates error processing (Brown & Braver, 2005). Behavioral work has furthermore shown that the effects of surprise and error processing interact, and that this interaction reflects a shared orienting response (Parmentier et al., 2019). In regards to inhibition, salient non-error events have indeed been shown to trigger non-selective CSE suppression (Wessel & Aron, 2013; Iacullo et al., 2020; Dutra et al., 2018; Tatz et al., 2021). As such, the hypothesis that errors invoke the same type of inhibition is straightforward, supporting the presence of a non-specific inhibition-orienting process after errors. Given its core role in the neural circuits underlying inhibition, it is tempting to hypothesize that this non-selective inhibition is due to the influence of the subthalamic nucleus, whose activity is correlated with non-selective CSE suppression (Wessel et al., 2016a), is active during the immediate post-error period (Siegert et al., 2014), and has non-selective effects on motor output (Parent & Hazrati, 1995; Mink, 1996).

Interestingly, the brain network underlying non-selective inhibition has recently been proposed to underlie even inhibitory effects on non-motor representations. Rapid, non-selective motor inhibition – like that observed shortly after error commission in the current study – is implemented via a hyper-direct pathway that involves broad excitatory projections from the subthalamic nucleus, which non-selectively activate the output nuclei of the basal ganglia. These nuclei, in turn, inhibit thalamocortical motor representations (Jahanashahi et al., 2015; Frank et al., 2007; Wessel & Aron, 2017). Notably, similar basal ganglia loops (including those involving thalamocortical projections that underpin the maintenance of active neural representations) also underlie other, non-motor processes (Redgrave et al., 2010; Guo et al., 2017; O’Reilly et al., 2014; Wei &Wang, 2016). Consequently, some more recent work has found that the circuits which govern the inhibition of motor activity may also underlie the inhibition of non-motor representations (Wessel et al., 2016b; Tempel et al., 2020; Apšvalka et al., 2022). Indeed, some have even found that the active inhibition of cognitive representations (e.g., associative memory) is accompanied by CSE reductions (Castiglione & Aron, 2021), further supporting the idea that motor and non-motor inhibition may recruit the same neural circuit. In the context of the current study, this is important because recent work has outlined potentially inhibitory effects of errors on non-motor, cognitive activity. For example, error commission leads to a suppression of the early sensory response to subsequent stimuli (Buzzell et al., 2017), slows the rate of evidence accumulation during active decision-making processes (Purcell & Kiani, 2016), and suppresses active working memory maintenance (Wessel et al., 2022b). It is possible that these are all common effects of the invocation of the non-selective, hyper-direct pathway for inhibitory control, one of whose key signatures is the CSE suppression found in the current study. While the current study conclusively demonstrates that errors do indeed trigger the non-selective inhibition of the motor system, the association between this inhibitory activity and the interruption of non-motor processes needs to be explicitly tested in future studies.

The second key finding of the current study is that motor excitability was selectively suppressed – i.e., at task-relevant muscles (Experiment 4) but not at task-irrelevant muscles (Experiment 3) – during the preparation of the next response following an error. Moreover, this suppression was predictive of post-error slowing, suggesting that this inhibitory process is more targeted and strategic than the non-selective suppression that takes place immediately after an error. This result was further buttressed by an exploratory re-analysis of the existing data from Wessel et al. (2019), in which we found that the neural generator underlying the fronto-central P3, functionally localized from a separate stop-signal task using ICA, was also active during the exact same post-error time-period and directly predicts PES. The stop-signal P3 purportedly reflects strategic, controlled adjustments to motor behavior after an initial inhibition-orienting phase (Diesburg & Wessel, 2021), similar to what has been purported to occur after errors by the AOT. Indeed, the stop-signal P3 is directly related to local inhibitory GABA_a_ activity in the motor system (Hynd et al., 2021). In the stop-signal task, the strategic, controlled adjustment comes in the form of the outright cancellation of the response, whereas during errors, it is a strategic slowing of its preparation in the service of higher accuracy (Danielmeier et al., 2011; King et al., 2010; Purcell & Kiani, 2016; Schiffler et al., 2017). As such, we propose that this mechanism strategically implements the selective inhibition of task-related effectors to purchase additional time for adaptive post-error adjustments to take effect.

Together, our study clearly demonstrates that errors are followed by the inhibition of the motor system. Moreover, inhibition is deployed in two different modes, which potentially map onto the purported two stages of post-error processing predicted by the adaptive orienting theory (Wessel et al., 2018; di Gregioro et al., 2018. Future work could causally test the association between inhibition and error processing by causally influencing the underlying neural network, e.g., via deep-brain stimulation of subthalamic nucleus (Cavanagh et al., 2011; Siegert et al., 2014; Wessel et al., 2016a; Zavala et al., 2016) and further explore the association between non-selective, broad inhibitory control and the non-motor effects of action errors.

## Acknowledgements

This work was funded by a grant from the National Institutes of Health (NIH R01 NS102201 to JRW). The authors would like to thank all the volunteers for their participation, Benjamin O. Rangel for code review, Teresa A. Treat for mixed effect model consultation, Joshua R. Tatz and Nathan Chalkley for pilot testing, and Maddie Carlson, Anna Kalan, Aarushi Dervesh, Meg Kester, and Ryan Potter for help in data acquisition.

